# Single-Particle Cryo-EM of Naturally Coexisting Mycoviruses Enables Structural Characterization of Conserved Capsid Folds, Divergent Architectures, and dsRNA Genome Organization

**DOI:** 10.64898/2026.07.07.737079

**Authors:** Guy Novoa, David Gil-Cantero, Juan M. Martínez-Romero, José R. Castón

**Author notes:** Correspondence: José R. Castón.

## Abstract

Mycoviruses frequently coexist within fungal hosts, yet high-resolution structural studies have traditionally relied on the purification of individual viral species, limiting the structural analysis of naturally occurring mixed infections. Here, we show that single-particle cryo-electron microscopy (cryo-EM) can simultaneously resolve multiple coexisting mycoviruses directly from naturally virus-infected fungal hosts. Using *Saccharomyces cerevisiae* strain TF229, which naturally harbours the closely related totiviruses Saccharomyces cerevisiae virus L-A (ScV-L-A) and L-BC (ScV-L-BC), we separated both viral populations *in silico* and reconstructed their capsids at near-atomic resolution under identical experimental conditions. Despite sharing only 10% sequence identity, their capsid proteins retain a highly conserved structural fold, whereas a ∼20° difference in asymmetric dimer orientation remodels capsid curvature and generates distinct virion architectures. Structural comparison further highlighted the C-terminal extension of ScV-L-BC, which mediates molecular swapping between neighbouring subunits and contributes to capsid stabilization. Symmetry relaxation revealed that, in both viruses, the encapsidated genome adopts a conserved spool-like organization, with ordered dsRNA filaments arranged into concentric layers. The outermost genome layer remains separated by ∼10–15 Å from the inner capsid surface, consistent with its predominantly electronegative character, while specific capsid–genome contacts are maintained mainly through the C-terminal regions of Gag. These findings reveal how conserved capsid folds can generate structurally distinct viral particles and establish mixed-sample cryo-EM as a scalable strategy for the high-resolution characterization of complex mycovirus communities.

**IMPORTANCE:** Fungal viruses are widabstacttespread and frequently occur as mixed infections, yet most structural studies rely on the purification of individual viruses, limiting our understanding of their diversity. This study demonstrates that cryo-electron microscopy can resolve closely related mycoviruses directly from naturally mixed samples, enabling their structural characterization under identical conditions. Application of this strategy to ScV-L-A and ScV-L-BC reveals how conserved capsid protein architectures can generate distinct viral particles and uncovers principles governing dsRNA genome organization. This work establishes a strategy to connect the rapidly expanding discovery of fungal viruses with structural and functional understanding.

## INTRODUCTION

Mycoviruses (fungal viruses) are widespread throughout the fungal kingdom, where multiple viral species frequently coexist within the same host, giving rise to complex virus–virus and virus–host interactions that influence viral persistence, transmission, and host phenotypes (1–4). Since the first description of a mycovirus in 1962, numerous species have been identified and characterized, revealing common biological features that include the general absence of an extracellular phase and transmission primarily through cell division, sporogenesis, and hyphal anastomosis (5–8). However, despite their widespread occurrence and biological importance, our current understanding of mycoviruses remains largely restricted to a relatively small number of fungal hosts of clinical, agricultural, and environmental relevance (6,9).

The widespread adoption of high-throughput sequencing (HTS) has transformed mycovirus research and dramatically increased the discovery of novel viral species from environmental and fungal samples worldwide (10). To date, several hundred mycovirus species assigned to 42 families have been reported, although their true diversity is thought to remain substantially underestimated (9,11). This methodological revolution has expanded the field beyond the inherent limitations of double-stranded RNA (dsRNA)-based detection methods and uncovered an unexpectedly broad genomic diversity that includes dsRNA, positive-sense single-stranded RNA [(+)ssRNA], negative-sense single-stranded RNA [(−)ssRNA], and single-stranded DNA (ssDNA) viruses (12–18). The expanding mycovirus virosphere exhibits remarkable diversity in viral lifestyles, genome organizations, and particle morphologies, as illustrated by members of the recently established families *Botourmiaviridae*, *Mymonaviridae*, and *Yadokariviridae* (19–22).

Despite this explosion in sequence-based virus discovery, high-resolution structural characterization has progressed much more slowly. To date, X-ray crystallography and cryo-electron microscopy (cryo-EM) structures are available for only a handful of mycoviruses, including Saccharomyces cerevisiae virus L-A (ScV-L-A), *Saccharomyces cerevisiae* virus L-BC (ScV-L-BC), Partitivirus stoloniferum virus F (PsV-F), Penicillium chrysogenum virus (PcV), Rosellinia necatrix quadrivirus 1 (RnQV1), and Rosellinia necatrix megabirnavirus 1 (RnMBV1) (23–28). Nevertheless, these studies have transformed our understanding of dsRNA virus particles. They demonstrate that viral capsids are not merely passive genome containers but dynamic molecular machines that harbour specialized functional elements. Examples include the cap-snatching active sites on the outer surfaces of ScV-L-A and ScV-L-BC (23,24), the arch domain of the PsV-F capsid protein (CP) involved in particle assembly and stability (25), and the putative proteolytic domain of the RnQV1 P2 protein (27). Comparative structural analyses further suggest that additional functional elements remain to be discovered within mycovirus capsids (29,30). However, the extension of high-resolution structural studies to the rapidly expanding mycovirus virosphere remains a major challenge because structural characterization still relies on the purification of individual viral species, an approach that is often hindered by low viral abundance and sample complexity (31)

The yeast *Saccharomyces cerevisiae* provides a well-established model for investigating naturally occurring mixed mycovirus infections. Across the diversity of *S. cerevisiae* strains, the dsRNA viruses Saccharomyces cerevisiae virus L-A (ScV-L-A) and Saccharomyces cerevisiae virus L-BC (ScV-L-BC), members of the family *Orthototiviridae*, are the most frequently detected and frequently coexist with the M dsRNA satellite viruses, which encode killer toxins that confer a competitive advantage to the host (32). Recent virome studies have further expanded our knowledge of the *S. cerevisiae* virome and have identified additional members of the families *Partitiviridae* and *Narnaviridae*, together with several less represented viral groups (33,34).

Among these viruses, ScV-L-A has historically served as the principal model for elucidating key molecular mechanisms, including ribosomal frameshifting and cap-snatching (23,35). Population-scale analyses have shown that ScV-L-A and ScV-L-BC frequently coexist within the same cells (36), where both viruses localize near polysomes to facilitate viral protein synthesis (37). Their long-term coexistence could reflect a complementary functional relationship: only ScV-L-A supports the replication and maintenance of M satellite viruses (32), whereas cap-snatching activity could also occur in trans between ScV-L-A and ScV-L-BC, which could increase the efficiency of viral transcription (38). Nevertheless, these chronic infections remain tightly controlled by host antiviral pathways, which prevent excessive viral replication that could become detrimental under environmental stress (39).

ScV-L-A and ScV-L-BC are closely related totiviruses that share a highly conserved structural and functional organization. Both viruses assemble ∼40 nm icosahedral capsids composed of 120 copies of the CP Gag, arranged as 60 asymmetric dimers that adopt two quasi-equivalent conformations (termed A and B) within a T = 1 icosahedral lattice (23,24,40). These capsids encapsidate a ∼4.6 kbp dsRNA genome with two open reading frames (ORFs), and viral translation produces either the Gag CP or a Gag–Pol fusion protein through programmed −1 ribosomal frameshifting, in which Pol corresponds to the viral RNA-dependent RNA polymerase (RdRP) (23,24,40). Despite this high degree of structural and functional conservation, the three-dimensional organization of the encapsidated genome and its relationship with the surrounding capsid remain largely unresolved.

To address the gap between mycovirus discovery and high-resolution structural characterization, we evaluated whether particle-classification strategies routinely used to resolve conformational and compositional heterogeneity could simultaneously separate and independently reconstruct distinct coexisting mycoviruses directly from a single mixed sample. As a proof of concept, we analyzed the naturally co-infecting ScV-L-A and ScV-L-BC virions present in *Saccharomyces cerevisiae* strain TF229, which allowed their direct structural comparison under identical experimental conditions. In addition, we applied symmetry relaxation to investigate the three-dimensional organization of their encapsidated dsRNA genomes, to gain new insights into the three-dimensional organization of encapsidated dsRNA in totiviruses.

## RESULTS

### Simultaneous cryo-EM reconstruction of ScV-L-A and ScV-L-BC from a single mixed sample

We analyzed a sample containing ScV-L-A and ScV-L-BC virions purified from *S. cerevisiae* strain TF229. SDS-PAGE analysis of virions purified from *S. cerevisiae* strain 1095 showed a major band, corresponding to the ScV-L-A CP (Gag) of 76 kDa (Fig. 1A, lane 1). In contrast, the TF229 sample displayed two major bands, corresponding to the CPs of ScV-L-A and ScV-L-BC of 76 and 81 kDa (Fig. 1A, lane 2). The ScV-L-BC Gag CP showed a faster electrophoretic mobility compared with ScV-L-A Gag, an anomalous electrophoretic mobility previously reported (38). Both samples also contained a minor band of a ∼180 kDa, corresponding to the Gag–Pol fusion protein of ScV-L-A, whereas the ScV-L-BC Gag–Pol protein was not detected because of its low abundance.

**Figure 1.**
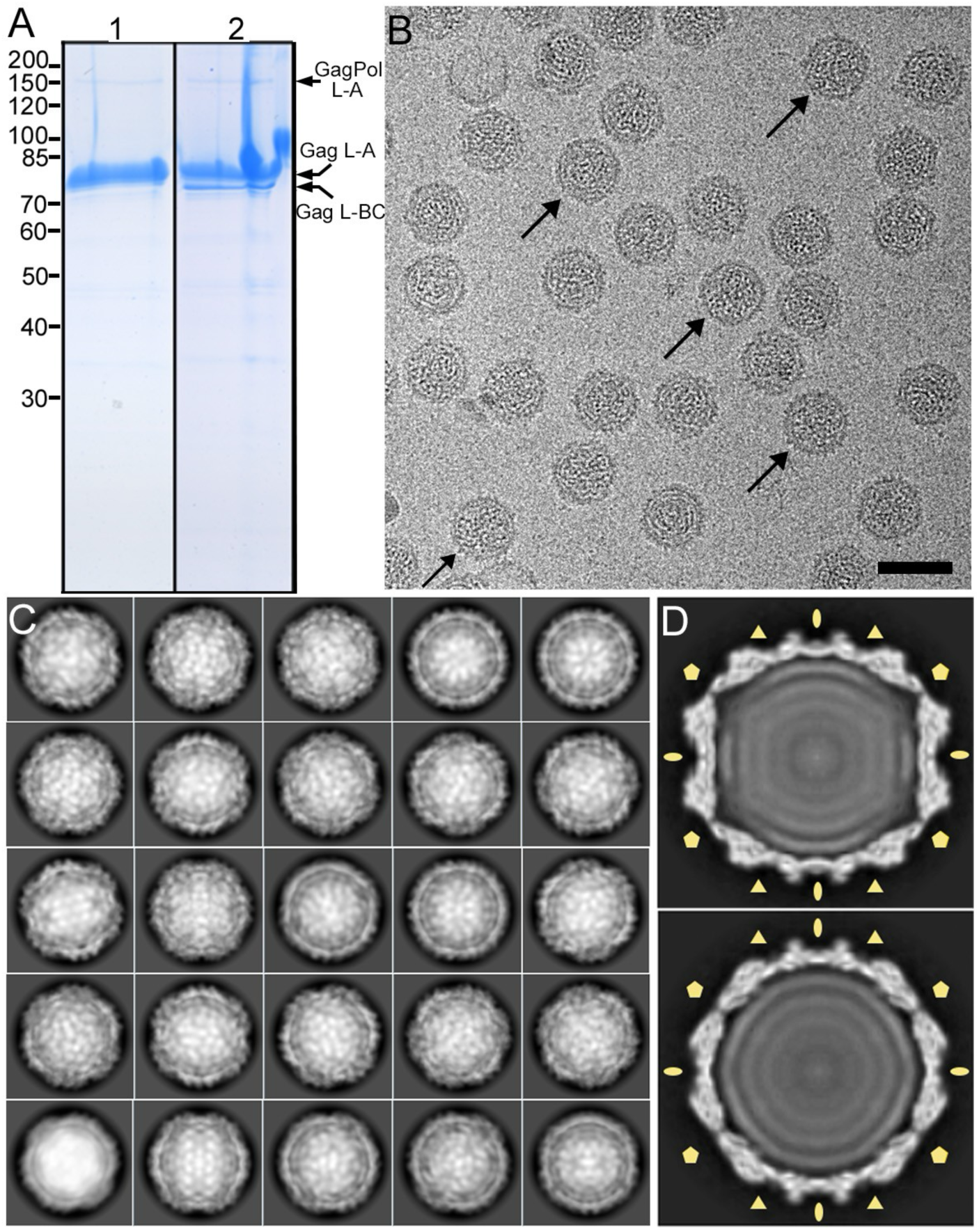
Biochemical and cryo-EM analysis of coexisting ScV-L-A and ScV-L-BC. (A) Coomassie blue-stained SDS-PAGE gels of virions purified from *S. cerevisiae*. Lane 1, virions purified from strain 1095 containing only ScV-L-A; lane 2, virions purified from strain TF229 containing ScV-L-A and ScV-L-BC. Arrows indicate the bands corresponding to the ScV-L-A and ScV-L-BC Gag capsid proteins and the ScV-L-A Gag–Pol fusion protein. Molecular size markers (kDa) are at left. (B) Cryo-EM micrograph of virions purified from strain TF229. Arrows indicate ScV-L-BC particles, identified after 3D classification and back-mapping of particle coordinates onto the original micrograph. Bar, 50 nm. (C) Representative reference-free 2D class averages of the extracted particles; ScV-L-A and ScV-L-BC could not be distinguished by visual inspection. (D) Representative central sections of the 3D reconstructions of ScV-L-A (top) and ScV-L-BC (bottom) after independent refinement, highlighting the polygonal morphology of ScV-L-A and the more rounded profile of ScV-L-BC. The icosahedral 2-, 3- and 5-fold symmetry axes are indicated by ovals, triangles and pentagons, respectively.

Cryo-EM analysis of the TF229 sample revealed an apparently homogeneous population of isometric particles composed of two different viral species, ScV-L-A and ScV-L-BC (Fig. 1B). A total of 469,633 particles were initially subjected to reference-free 2D classification. Visual inspection of the 2D classes did not allow discrimination between ScV-L-A and ScV-L-BC particles (Fig. 1C). However, subsequent 3D classification resolved two major particle populations corresponding to ScV-L-A (91% of particles) and ScV-L-BC (9%) when icosahedral symmetry was imposed (Fig. 1D). Despite their similar diameter of ∼42 nm, both capsids displayed clear structural differences in the 3D classes solved at ∼15–20 Å resolution (Fig. 1D). Central cross-sections revealed a polygonal profile in ScV-L-A (Fig. 1D, top) compared with the more rounded morphology of ScV-L-BC (Fig. 1D, bottom). The separation of ScV-L-A and ScV-L-BC by 3D classification allowed the assignment of both viral species in the original micrographs (Fig. 1B, arrows).

Both populations were independently refined with icosahedral symmetry that resulted in reconstructions of ScV-L-A and ScV-L-BC at 2.1 Å and 2.3 Å resolution, respectively, as estimated by the criterion of a 0.143 Fourier shell correlation (FSC) coefficient (Fig. S1). In agreement with previous structural studies (23,24,40), the high-resolution maps confirmed that both viruses have a T = 1 capsid formed by 120 copies of the CP Gag. These subunits are organized into 12 decamers located around the icosahedral 5-fold symmetry axes, with each decamer composed of five CP-A conformers and five intercalated CP-B conformers, which together define the 60 asymmetrical dimers forming the capsid shell (Fig. 2A–B; CP-A, blue; CP-B, yellow).

**Figure 2.**
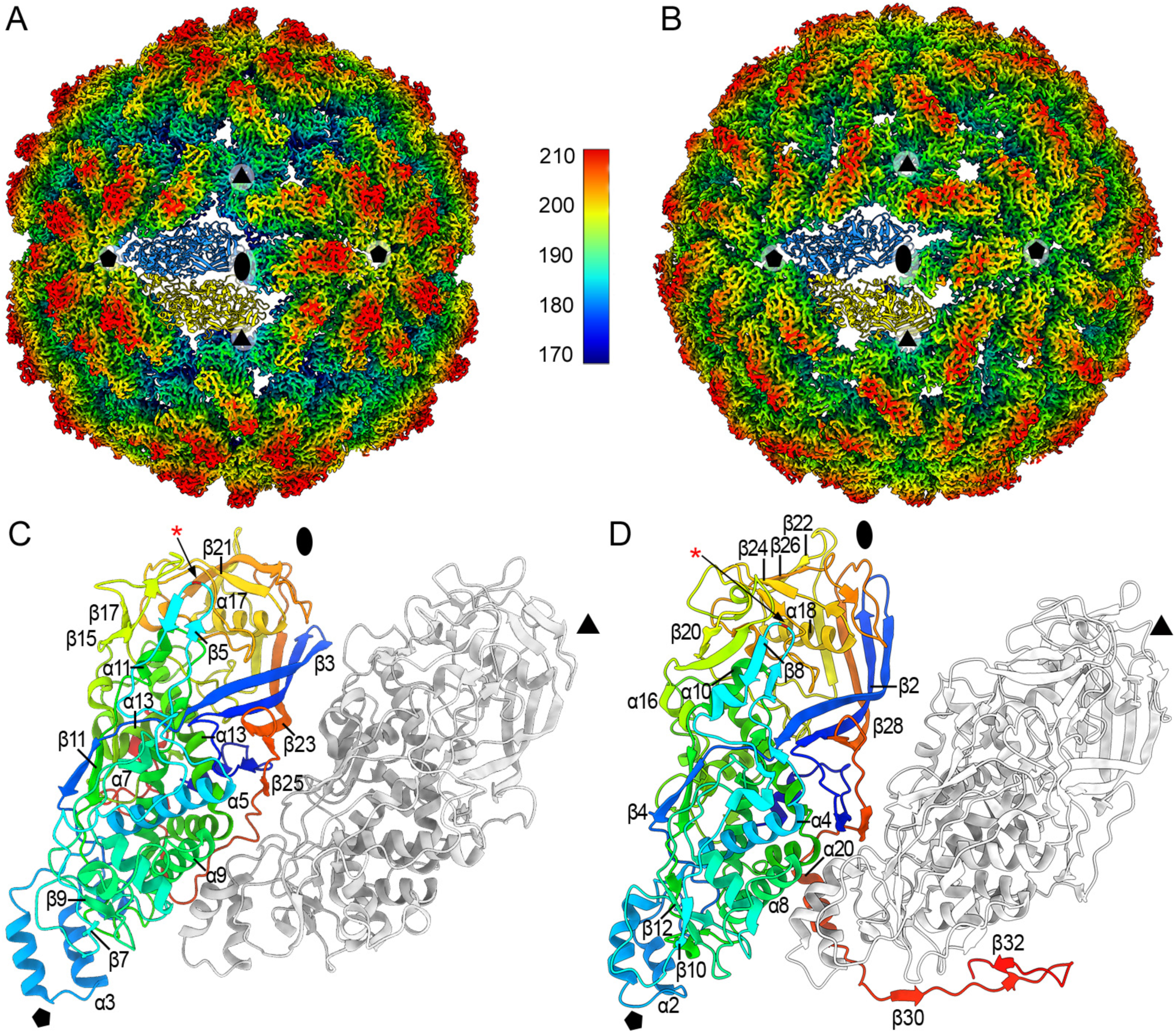
Three-dimensional cryo-EM reconstructions and capsid protein organization of ScV-L-A and ScV-L-BC. (A, B) Radially color-coded outer surfaces of ScV-L-A (A) and ScV-L-BC (B), viewed along a 2-fold axis of symmetry; the color key indicates the radial distance (in Å) from the particle center. The atomic structure of a CP dimer of subunits A (blue) and B (yellow) is shown. (C, D) Ribbon diagrams of the CP dimer of ScV-L-A (C) and ScV-L-BC (D) (top view). The A subunit is rainbow-colored from blue (N terminus) to red (C terminus); the B subunit is in gray. Secondary-structure elements are labelled and the icosahedral 2-, 3- and 5-fold symmetry axes are indicated by ovals, triangles and pentagons, respectively. The catalytic residues involved in cap-snatching activity (His154 in ScV-L-A and His156 in ScV-L-BC) are marked with a red asterisk.

### Atomic models of ScV-L-A and ScV-L-BC show conserved capsid protein folds

The atomic models obtained for ScV-L-A (PDB 9RK2) and ScV-L-BC (PDB 9SUJ) showed a high degree of structural conservation with previously reported structures (23,24). Superposition with the available ScV-L-A structure (PDB 1M1C) resulted in a root mean square deviation (RMSD) of 0.5 Å over 650 residues, whereas comparison with the ScV-L-BC structure (PDB 7QWZ) showed an RMSD of 0.5 Å over 647 residues.

In ScV-L-A, the Gag protein displays approximate dimensions of ∼100 × 35 × 50 Å in both quasi-equivalent conformations (Fig. 2C). A total of 653 out of 680 residues (Met1–Pro653) could be modelled, whereas the final 27 residues of the C-terminal region (Ala654–Glu680) remained unresolved in both conformers, most likely due to their intrinsic flexibility or structural disorder. Both CP-A and CP-B contain 18 α-helices and 28 β-strands, with the visible N- and C-terminal regions oriented toward the capsid interior (Fig. S2).

The catalytic residue involved in cap-snatching activity, His154, is located within β-strand 5 in the protruding region of the capsid protein (Fig. 2C, red asterisk). This residue adopts an exposed orientation compatible with the formation of the transient covalent intermediate required for the transfer of the mRNA cap to newly synthesized viral transcripts (23,41).

In ScV-L-BC, the Gag protein exhibits similar dimensions, although the C-terminal region of CP-A extends ∼50 Å beyond the compact core of the monomer (Fig. 2D). The final model contains 658 of the 697 residues in CP-A (Met1–Pro658) and 648 residues in CP-B (Met1–Thr620 and Arg631–Pro658). The unresolved internal region in CP-B (Gly621–Leu630) corresponds to the segment that forms α-helix 20 in CP-A, which indicates increased flexibility of this region in the B conformer. Similar to ScV-L-A, the visible N- and C-terminal regions of ScV-L-BC are oriented toward the capsid interior. The ScV-L-BC CP contains 20 α-helices in CP-A (19 in CP-B) and 32 β-strands (Fig. S3).

In contrast to ScV-L-A, ScV-L-BC exhibits β-strand complementation between neighbouring subunits through a β-augmentation mechanism (Fig. S4A, B). In the CP-A conformer, β-strand 30 extends into the adjacent CP-A subunit within the same pentamer and becomes incorporated between β-strands 16 and 17, which establishes an intermolecular exchange between equivalent subunits through molecular swapping (Fig. S4C). In contrast, although CP-B also forms a β16–β17–β30 sheet, this interaction occurs intramolecularly, with all three β-strands belonging to the same capsid protein (Fig. S4D-F).

The catalytic residue His156, which is equivalent to His154 in ScV-L-A, occupies a conserved position within the protruding region of the capsid protein. Thus, the catalytic histidine residues involved in cap-snatching activity occupy equivalent positions in both viral capsids (Fig. 2D, red asterisk).

Despite sharing only 10% sequence identity (Fig. 3A, residues highlighted in red) and 22% sequence similarity (Fig. 3A, residues boxed in blue), ScV-L-A and ScV-L-BC CPs exhibit a remarkable degree of structural conservation. Superposition of both Gag proteins resulted in RMSD values of 1.1 Å and 1.3 Å for the aligned residues in the A and B conformers, respectively. This structural conservation suggests a strong evolutionary pressure to preserve key elements involved in capsid protein folding and/or capsid assembly. The conserved structural core includes 15 α-helices and 20 β-strands out of a total of 18–20 α-helices and 28–32 β-strands present in both proteins (Fig. 3A).

**Figure 3.**
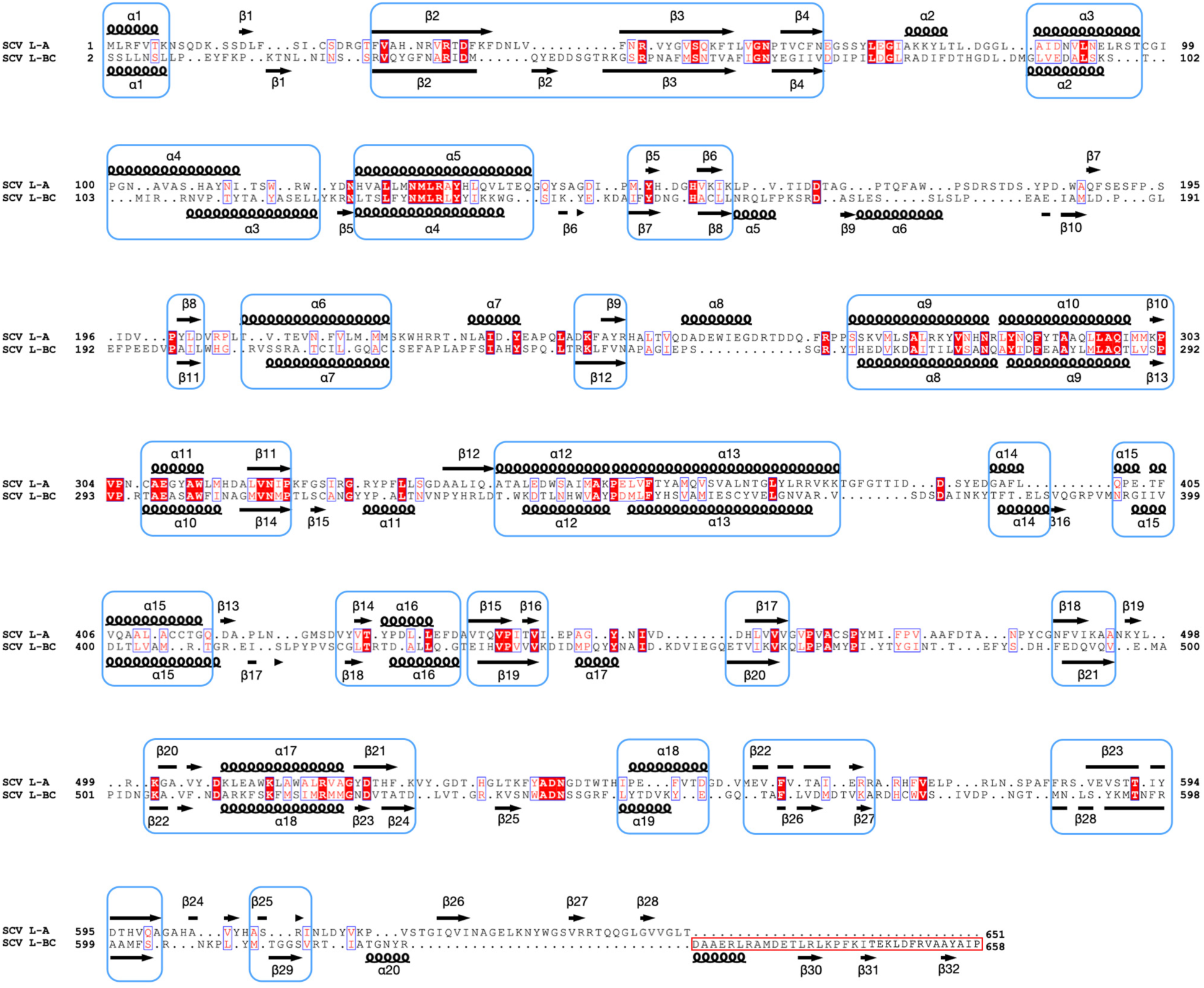
Structure-based sequence comparison of ScV-L-A and ScV-L-BC capsid proteins. Structure-based sequence alignment of the ScV-L-A and ScV-L-BC capsid proteins in conformation A. Identical residues, white letters on a red background; partially conserved residues, red. Gaps in the alignment are represented by dots. Secondary-structure elements are indicated above (ScV-L-A) and below (ScV-L-BC) the sequence alignment (spirals, a-helices; arrows, b-strands). Conserved SSE are highlighted in blue rectangles. The C-terminal region from A subunits involved in molecular swapping is indicated (red rectangle).

### Differences in capsid morphology arise from changes in Gag protein orientation

Despite the high structural conservation between ScV-L-A and ScV-L-BC Gag proteins, their assembly results in capsids with clearly distinct morphologies. Central sections of the atomic models showed that these differences mainly arise from variations in the relative orientation and tilt of the capsid protein subunits (Fig. 4A, B).

**Figure 4.**
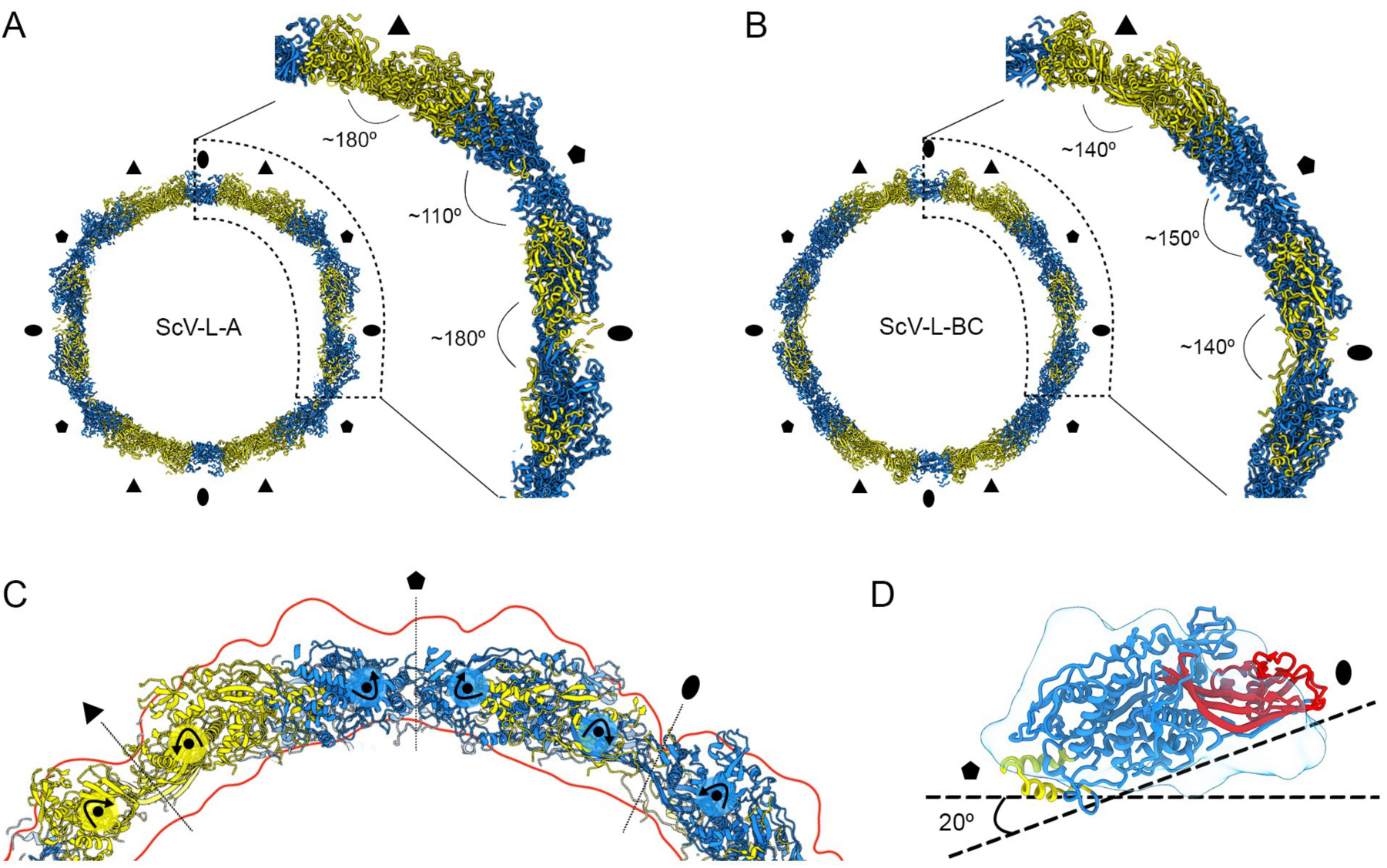
Differences in capsid morphology arise from changes in Gag protein orientation. (A, B) Central sections of the atomic models of the ScV-L-A (A) and ScV-L-BC (B) capsids. CP-A and CP-B conformers are shown in blue and yellow, respectively. Regions containing the 2-, 3- and 5-fold symmetry axes (ovals, triangles and pentagons) are enlarged on the right, with the intersubunit angles indicated. (C) Superposition of the ScV-L-BC atomic model onto the ScV-L-A capsid outline (red), showing the relative rotation of the capsid proteins around their central region. (D) Atomic model of the ScV-L-BC capsid protein in conformation A, colored according to structural regions (region 1, yellow; region 2, blue; region 3, red) and superimposed onto the ScV-L-A CP-A outline (blue). The ∼20° change in tilt that would be required for the ScV-L-BC capsid protein to adopt an ScV-L-A-like orientation is indicated.

Analysis of the angles formed between neighbouring Gag proteins at the different symmetry axes explains the structural basis of these morphological differences. Around the 2- and 3-fold symmetry axes, ScV-L-A subunits adopt an almost coplanar arrangement, with intersubunit angles close to 180°. In contrast, ScV-L-BC subunits display a more curved organization, with angles of approximately 140°. The opposite behaviour is observed around the 5-fold symmetry axes, where ScV-L-A subunits form a sharper angle of approximately 110°, whereas ScV-L-BC adopts a wider angle of approximately 150°. Therefore, the ScV-L-A capsid concentrates curvature in specific regions through the combination of highly curved (∼110°) and nearly planar (∼180°) interfaces, whereas the intermediate angles observed in ScV-L-BC (∼140–150°) generate a more regular spherical architecture (Fig. 4A, B; Movie S1).

The transition from the ScV-L-BC architecture (Fig. 4C, atomic model) to the ScV-L-A architecture (Fig. 4C, red contour) requires an ∼20° rotation of each asymmetric unit around an axis perpendicular to the intradimer interface. This rotation affects both CP-A and CP-B conformers and modifies the relative position of the CP domains.

To describe the structural rearrangements responsible for these morphological differences, each Gag subunit was divided into three structural regions: region 1, composed mainly of helices α2 and α3 located near the 5-fold symmetry axis (Fig. 4D, yellow); region 2, which contains most secondary structure elements and defines the rotation centre (Fig. 4D, blue); and region 3, a β-strand-rich region that mediates interactions with neighbouring subunits across the 2- and 3-fold symmetry axes (Fig. 4D, red). The ∼20° rotation of each asymmetric dimer shifts region 1 toward the particle interior in ScV-L-BC, whereas region 3 becomes elevated relative to ScV-L-A. Despite this global rearrangement, the catalytic residues His154 in ScV-L-A and His156 in ScV-L-BC remain oriented toward the groove formed between regions 2 and 3, preserving the structural environment required for cap-snatching activity. These structural rearrangements explain how highly conserved capsid proteins generate distinguishable capsid architectures while maintaining conserved functional sites.

### Capsid–genome interactions in ScV-L-A and ScV-L-BC

The 3D icosahedrally averaged reconstructions of full ScV-L-A and ScV-L-BC particles showed internal densities corresponding to the packaged dsRNA genomes. These densities appeared as two to three concentric layers beneath the capsid shell, without detailed structural features and became progressively more diffuse toward the centre of the particle. Analysis of the unsharpened maps of the full virions showed CP–RNA contacts in specific regions of both ScV-L-A (Fig. 5A) and ScV-L-BC (Fig. 5B), mainly mediated by the CP ordered C-terminal regions. In the CP-B conformers, the interaction occurs underneath the corresponding capsid protein in both ScV-L-A and ScV-L-BC (Fig. 5C, D; green). In contrast, although the C-terminal regions of CP-A are oriented toward the genome in both viruses, this connection is weaker in ScV-L-A (Fig. 5C, left; red). In ScV-L-BC the interaction is established through the C-terminal region involved in molecular swapping (Fig. 4D, left; red).

**Figure 5.**
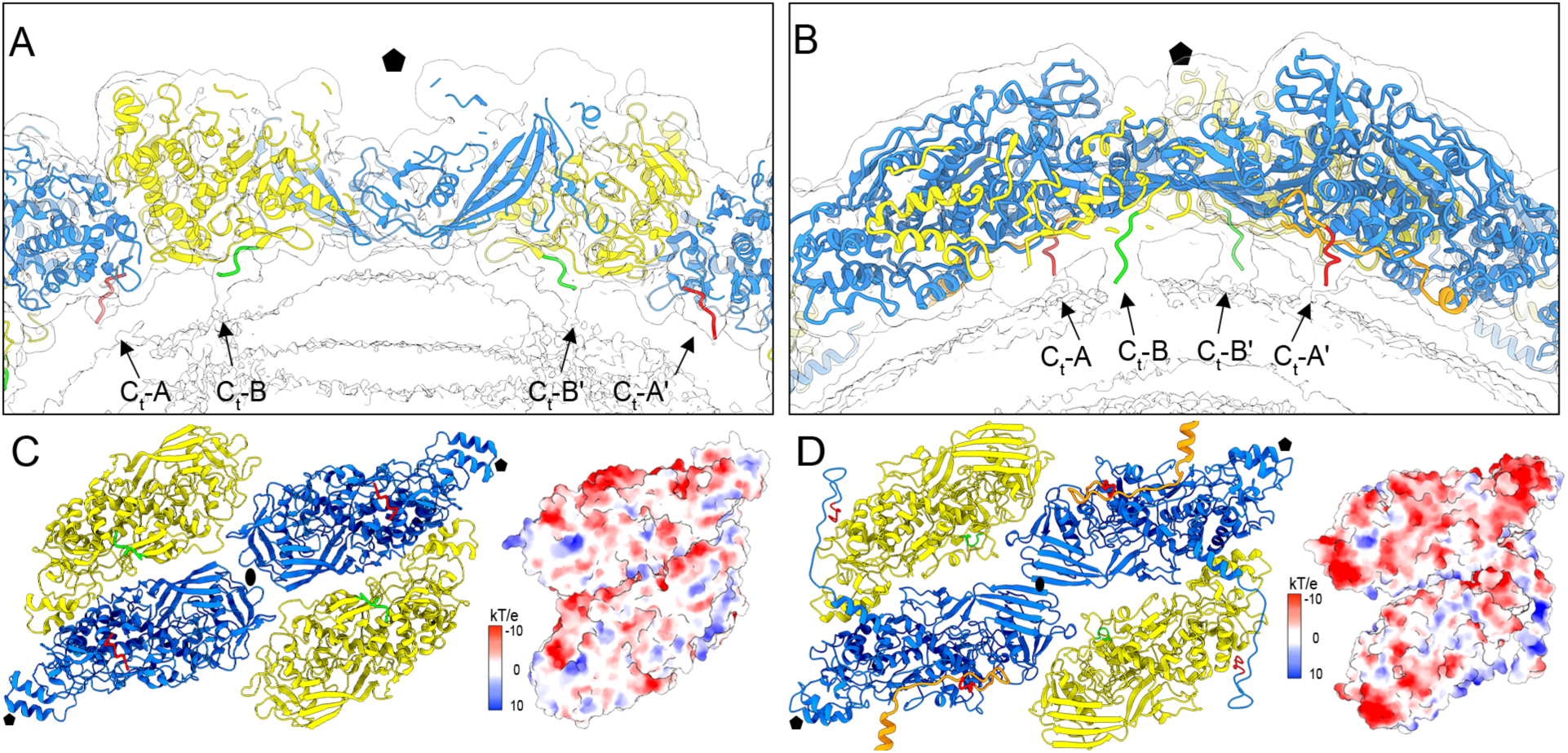
Capsid–genome interactions and electrostatic properties in ScV-L-A and ScV-L-BC. (A, B) Central sections of unsharpened cryo-EM maps of full ScV-L-A (A) and ScV-L-BC (B) particles contoured at 1s above the mean density, showing capsid–genome interactions preserved after icosahedral averaging. Detailed views around a 5-fold symmetry axis (pentagon) are shown, with the two capsid protein conformers, CP-A (blue) and CP-B (yellow), and the outermost dsRNA genome layer (white). The C-terminal regions involved in capsid–genome contacts are labelled (Ct-A, Ct-B, Ct-B′, and Ct-A′). (C, D) Inner view of a tetramer of ScV-L-A (C) and ScV-L-BC (D) CPs formed around the 2-fold symmetry axis (oval) (left). In ScV-L-A, the C-terminal regions of the A and B subunits are shown in red and green, respectively. In ScV-L-BC, the C-terminal regions of the B subunits are shown in green, whereas two C-terminal regions from adjacent A subunits involved in molecular swapping with neighbouring A subunits are highlighted in orange (left). Views of the dimer inner surfaces represented with electrostatic potentials showing the distribution of negative (red) and positive (blue) charges (right).

To further characterize the capsid–genome interface, the electrostatic potential of the complete capsid was calculated. In ScV-L-A, the inner capsid surface contained both positive and negative charges, with ∼20% and 6% of the asymmetric unit surface corresponding to negatively and positively charged regions, respectively (Fig. 5A, right). The inner surface of ScV-L-BC displayed a stronger electronegative character than ScV-L-A, with negatively and positively charged regions accounting for ∼33% and 5% of the asymmetric unit surface, respectively (Fig. 5D, right).

Despite the contacts observed between the capsid inner surface and the outermost genome layer, the dsRNA remains separated from the capsid shell by approximately 10–15 Å in both ScV-L-A and ScV-L-BC. This separation suggests that the genome retains a certain degree of mobility, which could be favoured by the predominantly electronegative character of the inner capsid surface.

### Symmetry relaxation reveals the internal organization of the dsRNA genome

The organization of the packaged genomes in ScV-L-A and ScV-L-BC was further investigated without imposing the icosahedral symmetry of the capsid. To this end, the icosahedrally ordered component corresponding to the capsid shell was subtracted from the original particles, and the remaining internal densities were analyzed by 3D classification with symmetry relaxation. This approach allowed the asymmetric components inside the capsid to be reconstructed independently of the global icosahedral symmetry and revealed whether equivalent genome configurations were present among different particles.

For ScV-L-A, symmetry relaxation was performed using 280,555 particles. A spherical mask with a radius of 180 Å was applied during capsid subtraction and 3D classification. The resulting classes contained 33, 26, 21 and 20% of the particles and showed similar genome organizations, with dsRNA arranged as a spool-like structure composed of three to four layers of parallel filaments surrounding a disordered central region (Fig. 6A). Although the filament arrangement differed between classes, the overall organization was conserved. The resolved filaments showed a diameter of ∼20 Å, consistent with A-form dsRNA (∼23 Å diameter), as confirmed by fitting an atomic model of dsRNA into the density (Fig. 6B). Some filaments displayed continuity between different genome layers, including two filaments located near the equatorial region that connected layer 1 with layer 2 and vice versa (Fig. 6B, arrows). The distance between neighbouring filaments was approximately 35 Å, both within the same layer and between adjacent layers (Fig. 6C). Based on an axial rise of 2.81 Å/base pair for A-form dsRNA and a genome size of 4.6 kbp, the expected genome length is approximately 12,926 Å. The total length of the traced filaments in the symmetry-relaxed reconstruction was approximately 9,000 Å, corresponding to ∼70% of the ScV-L-A genome.

**Figure 6.**
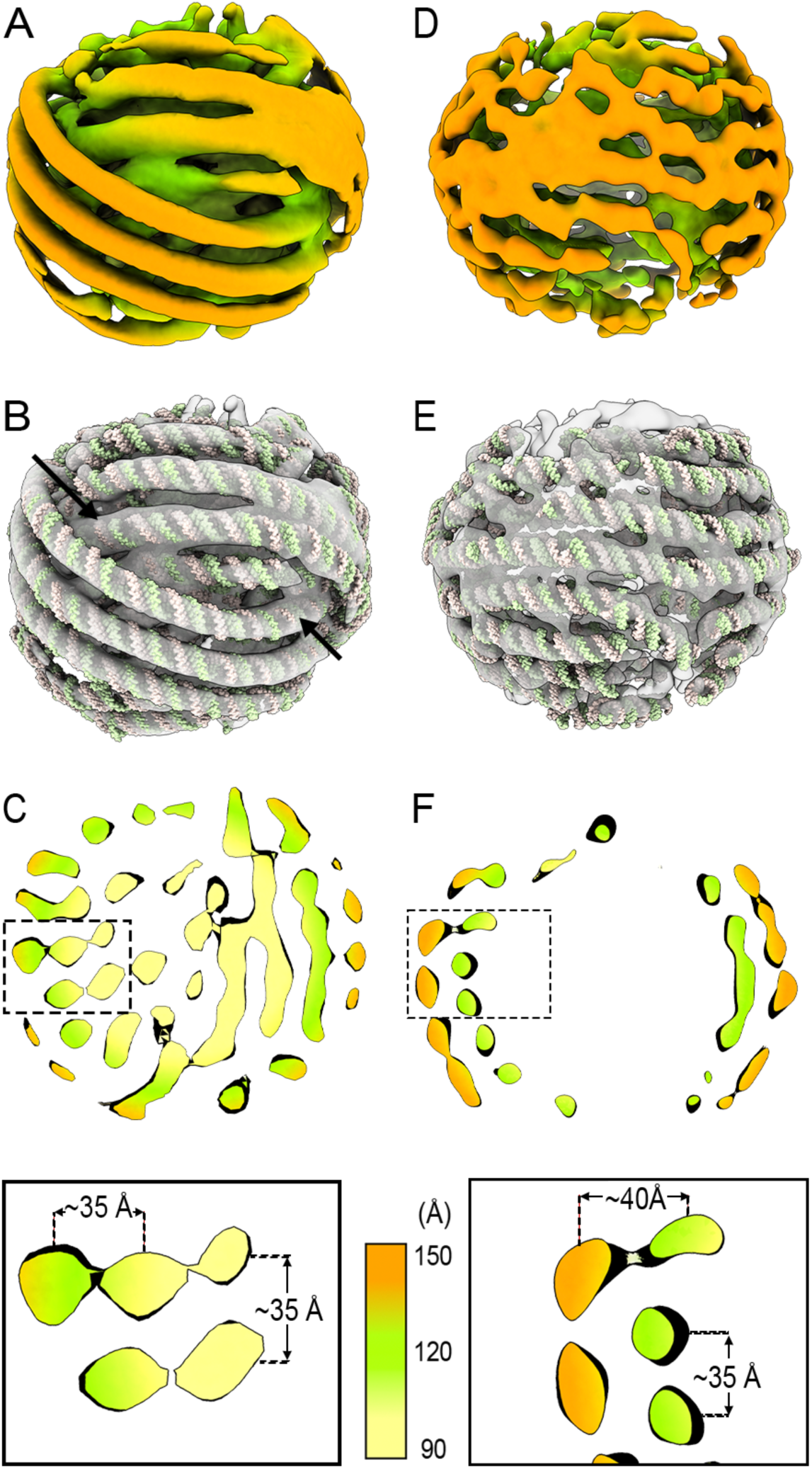
Three-dimensional organization of the packaged dsRNA genome resolved by symmetry relaxation. (A–C) ScV-L-A; (D–F) ScV-L-BC. (A, D) External views of the ordered genome densities colored according to radial distance from the particle center (Å; color scale in C), showing the three-dimensional arrangement of dsRNA filaments within the capsid. (B, E) Segmented representation of the genome densities, illustrating the spool-like organization of the dsRNA filaments. In ScV-L-A, black arrows indicate continuous genome filaments transitioning between adjacent RNA layers. (C, F) Central sections of the genome densities colored according to radial distance. Boxed regions are enlarged below, showing the spacing between adjacent dsRNA filaments (∼35 Å in ScV-L-A; ∼35–40 Å in ScV-L-BC).

The same strategy was applied to ScV-L-BC using 29,699 particles, except that we applied an annular mask with external and internal radii of 184 Å and 110 Å, respectively. The resulting classes contained 42, 27, 21 and 10% of the particles (Fig. 6D). The ScV-L-BC genome displayed a similar spool-like organization to that observed in ScV-L-A, although filament discontinuities limited complete tracing (Fig. 6E). The resolved filaments also showed a diameter of approximately 20 Å and were separated by 35–40 Å (Fig. 6F). Because the shell mask excluded the innermost genome region, the total length of the traced filaments was limited to approximately 7,000 Å, corresponding to ∼55% of the 4.6 kbp ScV-L-BC genome.

## DISCUSSION

The 120-subunit T=1 capsids of mycoviruses and related dsRNA viruses belonging to the Bluetongue virus (BTV)-like lineage exhibit different organizational strategies depending on the configuration of their asymmetric units. Based on currently available structures, four main architectural classes can be distinguished. The first class corresponds to symmetric homodimeric capsids, as observed in the partitivirus PsV-F (25), where two structurally equivalent CP subunits form a symmetric dimer around a local 2-fold symmetry axis, similar to the organization described in picobirnaviruses (42). The second class includes pseudo-T = 2 (P = 2) capsids, such as RnQV1 (27), whose asymmetric unit consists of two different but structurally related CPs.

The most common organization corresponds to asymmetric homodimeric capsids, where a single CP adopts two quasi-equivalent conformations within the asymmetric unit. This architecture is shared by the mycoviruses analyzed in this study (ScV-L-A and ScV-L-BC), as well as several protozoan totiviruses, including Giardia lamblia virus (29), Leishmania RNA virus 1 (43), and Trichomonas vaginalis virus 2 (44). Finally, a fourth class consists of asymmetric homodimeric capsids that incorporate additional structural proteins, as described for RnMBV1 (28) and the animal totivirus Omono River virus (OmRV) (45), where accessory proteins decorate specific symmetry axes. Although several lower-resolution structures lack atomic models, their capsid organization suggests that they also conform to these architectural categories, including HvV190S (46) and infectious myonecrosis virus (IMNV) (47). The presence of additional external domains in extracellularly transmitted totiviruses, such as OmRV and IMNV, may reflect adaptations associated with receptor recognition and cell entry, whereas most mycoviruses with strictly intracellular transmission generally lack these protruding elements.

In this study, we demonstrate that modern cryo-EM image processing workflows can resolve the structures of naturally coexisting mycoviruses directly from a single biological sample. Using ScV-L-A and ScV-L-BC as a model system, we show that classification-based approaches can separate highly similar viral populations that cannot be distinguished by visual inspection of cryo-EM images or 2D classes. Although this ability could be anticipated from the capacity of current image-processing algorithms to resolve subtle conformational and compositional heterogeneity, the separation of two closely related viral species demonstrates the potential of *in silico* purification strategies for the structural characterization of mixed viral infections. The resulting near-atomic resolution reconstructions are consistent with previously determined structures (23,24), and confirm that mixed viral populations can be exploited for simultaneous structural characterization. This strategy provides an alternative to labor-intensive purification of individual viral species and may facilitate the analysis of complex mycovirus communities, which are frequently found in fungal hosts.

Despite sharing only limited sequence identity, ScV-L-A and ScV-L-BC capsid proteins preserve a highly conserved structural fold. However, their assembly results in clearly distinct capsid morphologies due to differences in the relative orientation of the asymmetric dimers. A ∼20° variation in dimer tilt modifies the distribution of capsid curvature, which generates the more angular morphology of ScV-L-A and the smoother, more spherical architecture of ScV-L-BC. These structural rearrangements also modify the dimensions of the 2-, 3-, and 5-fold pores, among which the 5-fold channel represents the largest opening in both viruses and is substantially expanded in ScV-L-A. Therefore, small changes in the relative orientation of conserved structural units can be amplified during assembly to generate different capsid architectures.

In addition to differences in capsid protein orientation, ScV-L-BC contains a specific C-terminal extension that contributes to capsid stabilization through molecular swapping. The C-terminal region of CP-A (Thr620–Pro658), which is involved in the exchange between neighbouring subunits, is responsible for the increased number of intermolecular interactions observed in ScV-L-BC. Computational removal of this region reduces the number of intersubunit contacts to values comparable to those observed in ScV-L-A, highlighting its role as a structural reinforcement element. This strategy, mediated by the ∼35-residue C-terminal extension of ScV-L-BC CP-A, is also observed in other dsRNA viruses, where terminal extensions strengthen protein–protein interfaces. Similar examples include the C-terminal extension of PcV P2 (∼25 residues) (26), the C-terminal regions of RnQV1 P2 (∼95 residues) and P4 (∼15 residues) (27), the C-terminal extensions of RnMBV1 CP-A and CP-B (∼50 residues) (28), and OmRV CP-A (∼35 residues) and CP-B (∼5 residues) (45). These examples suggest that terminal extensions represent a recurrent strategy among dsRNA viruses to reinforce capsid assemblies while maintaining a conserved capsid protein fold. Consistent with the enhanced stability associated with the ScV-L-BC capsid, ScV-L-BC VLPs have been proposed as promising platforms for future biotechnological applications (48).

Our cryo-EM analysis showed a clear predominance of ScV-L-A particles (∼91%) over ScV-L-BC particles (∼9%) in the naturally co-infected *S. cerevisiae* TF229 strain. This ratio is consistent with recent analyses performed from cellular extracts, which reported a similar ScV-L-A/ScV-L-BC ratio (37). In addition, these studies showed that ScV-L-A particles accumulate in specific cellular regions, where they associate with ribosomes and form polysome-associated viral clusters that promote immediate translation of viral transcripts while protecting viral mRNAs from degradation (37). The stable coexistence of both viruses despite their different abundance suggests that chronic mixed infections are maintained through a complex balance between viral replication, host regulation, and potential functional interactions between viral species.

Although ScV-L-A and ScV-L-BC can complete their replication cycles independently, their frequent co-occurrence in *S. cerevisiae* populations suggests that mixed infections may provide selective advantages under specific conditions. ScV-L-A has traditionally been considered a persistent virus with limited impact on host viability; however, excessive viral replication can become detrimental in strains with impaired antiviral pathways due to proteotoxic stress (39). Therefore, controlled maintenance of ScV-L-A might reflect an equilibrium between viral persistence and host tolerance.

In this context, the structural analyses performed in this study, together with previous physicochemical characterization (48), support the high resistance of the ScV-L-BC capsid to physical and chemical stress. This enhanced stability, combined with the ability of both ScV-L-A and ScV-L-BC to act as donors and acceptors of 5′ mRNA caps during cap-snatching reactions, may provide a reservoir of functional viral particles capable of maintaining transcriptional activity even under unfavourable conditions. Therefore, the increased structural stability of ScV-L-BC could contribute to its persistence within mixed viral populations. This coexistence indirectly supports the maintenance of M satellite viruses, which depend specifically on ScV-L-A replication and confer the killer phenotype that provides a competitive advantage against other *S. cerevisiae* strains (38,39).

The organization of the encapsidated dsRNA genomes appears to be strongly influenced by the physicochemical properties of the capsid interior. In both ScV-L-A and ScV-L-BC, the genome is organized into concentric dsRNA layers that remain separated from the inner capsid surface by ∼10–15 Å. This separation is consistent with the predominantly electronegative character of the capsid interior, which likely prevents extensive nonspecific RNA–protein interactions. The strongest CP–RNA contacts observed in ScV-L-A and ScV-L-BC are mediated by the C-terminal regions of the CPs and involve the outermost genome layers. Together with the progressive loss of ordered density toward the particle core, these observations support a model in which the external dsRNA layers are more constrained by capsid interactions, whereas internal genome regions remain more flexible.

Symmetry relaxation provided further insight into the asymmetric organization of the packaged genome. In both viruses, dsRNA filaments adopt a spool-like arrangement, although this organization was better resolved in ScV-L-A due to the larger number of particles and higher quality of the reconstruction. This spool-like organization suggests that the packaged dsRNA is not randomly distributed within the capsid but instead adopts recurrent configurations that are partially coupled to the icosahedral shell.

Beyond the structural insights into ScV-L-A and ScV-L-BC, this work establishes mixed-sample cryo-EM as a scalable strategy for the characterization of naturally coexisting mycoviruses. This approach reduces the experimental limitations associated with individual virus purification and provides an opportunity to investigate viral communities in a more native biological context, where multiple viral species may coexist and establish long-term interactions with each other and their fungal hosts. As high-throughput sequencing continues to reveal increasingly complex fungal viromes, scalable structural approaches will be essential to connect viral discovery with biological function.

## MATERIALS AND METHODS

### Yeast strains, viruses and virion purification

*Saccharomyces cerevisiae* strain TF229, which naturally harbours both ScV-L-A and ScV-L-BC, was used for the purification of both viruses, whereas strain 1095, which contains only ScV-L-A, was used for the purification of ScV-L-A alone. The same purification protocol was applied to both strains.

Yeast cultures were grown at 30 °C for 4 days under gentle agitation in YPAD medium (1% yeast extract, 2% peptone, 0.04% adenine sulphate, 2% potato dextrose, 94.96% water). Cells were harvested by centrifugation at 9,000 x g for 5 min and washed with distilled water. The resulting wet biomass was approximately 80 g for each strain.

Yeast cells were resuspended in lysis buffer (100 mM Tris pH 7.6, 1 M sorbitol, 20 mM β-mercaptoethanol, and 0.5 mg zymolyase) and incubated for 1 h at 37 °C. The suspension was centrifuged at 3,500 x g for 5 min, and the resulting pellet was resuspended in buffer A (50 mM Tris-HCl pH 7.8, 150 mM NaCl, 5 mM EDTA) supplemented with 1 mM DTT. Cell lysis was performed with a *French* press at 1000 bars. The homogenate was clarified by centrifugation at 9,000 x g for 20 min, and the supernatant was further centrifuged at 130,000 x g for 40 min. The resulting pellet was resuspended in 3 ml of buffer A and loaded onto a 12-mL CsCl solution (initial density of 1.35 g/cm^3^), which generated a self-generated density gradient during centrifugation at 215,000 x g for 13 h.

Fractions of 1 ml were collected from the top of the tube. Following SDS-PAGE analysis, the fraction containing virions was diluted in 7 volumes of buffer A and concentrated to ∼50 μl using Amicon centrifugal filters (Millipore) at ∼14,000 x g for 20 min.

### Cryo-EM specimen preparation and data collection

Samples were vitrified using a Vitrobot Mark IV (FEI) (22 °C, 95% humidity, blot force -3, blot time 3 s). A 4-μL aliquot of sample was applied to a glow-discharged grid. Carbon-coated Cu/Rh grids (Quantifoil) were used and stored in grid containers in liquid nitrogen until screening. Data were acquired on a Titan Krios (Thermo Fisher Scientific) transmission electron microscope operated at 300 kV, equipped with a Gatan K3 direct electron detector operating in super-resolution mode. Movies consisted of 30 frames, with a total dose of 31.2 e^-^/Å^2^. The nominal magnification was 81,000x, corresponding to a calibrated pixel size of 1.06 Å. The defocus range for the dataset was from -0.8 to -2.3 μm, with a 0.3 μm step. Automated acquisition was performed using EPU (Thermo Fisher Scientific), resulting in 12,248 images. A summary of the data collection and refinement statistics is provided in Table S1.

### Image processing

Movie frames were aligned and dose-weighted using MotionCor2, and contrast transfer function (CTF) parameters were estimated with CTFFIND4 (49). Dose-weighted micrographs were subjected to automated particle picking with Xmipp3 (50), yielding a total of 469,633 particles. Particles were extracted using a box size of 480 pixels and subjected to reference-free 2D classification to remove damaged particles and contaminants. The selected particle set was subsequently classified into five 3D classes, four corresponded to ScV-L-A and one to ScV-L-BC. Two ScV-L-A classes were merged and, together with the single ScV-L-BC class, were independently refined in cryoSPARC imposing icosahedral symmetry (51). The overall resolution of each reconstruction was estimated according to the gold-standard Fourier shell correlation (FSC = 0.143) criterion, and local resolution was estimated as implemented in Scipion using MonoRes (52).

### Model building, refinement and validation

Atomic models were built and refined for the asymmetric units of ScV-L-A and ScV-L-BC. Using as initial models the X-ray and cryo-EM structures of ScV-L-A (PDB 1M1C) and ScV-L-BC (PDB 7QWZ), respectively, these were fitted as rigid bodies into the corresponding cryo-EM density maps, and subjected to iterative real-space refinement using Phenix (53), with global minimization, local grid search and atomic displacement parameter (B-factor) refinement enabled, alternated with manual adjustment in Coot (54). The quality of the refined models was validated with MolProbity, as implemented in Phenix (55) and with the World-wide PDB (wwPDB) OneDep System (https://deposit-pdbe.wwpdb.org/deposition). Refinement and validation statistics are summarized in Table S1.

### Genome organization by symmetry relaxation, electrostatic analysis and structural comparison

To reconstruct the internal organization of the packaged genomes, the icosahedral symmetry was relaxed. Icosahedral alignment parameters were used as the initial reference, and particle orientations were locally refined without symmetry constraints using a mask encompassing the region that includes the genome density. This approach enabled visualization of the ordered dsRNA layers within the viral capsids.

The approximate number of base pairs contained within each resolved RNA layer was estimated from the traced filament length and the axial rise of A-form dsRNA. The electrostatic potential of the inner capsid surface was calculated from the atomic models using ChimeraX (56), and the relative proportions of positively and negatively charged regions were estimated.

Pore diameters at the 2-, 3-, and 5-fold symmetry axes were measured using ChimeraX. Structural superpositions, root-mean-square deviation (RMSD) calculations, and structure-based sequence comparisons were performed using the Dali server (57).

## Data availability

The atomic coordinates and cryo-EM density maps were deposited in the Protein Data Bank (www.pdb.org) and EM Data Bank (www.emdatabank.org) with codes 9RK2 and EMD-54013 for the ScV-L-A virions, and 9SUJ and EMD-55221 for the ScV-L-BC virions.

## Author contributions

J.R.C. conceived and designed the study. G.N., D.G.-C. and J.M.M.-R. performed the experiments and analysed the data. J.R.C. supervised the work and acquired funding. G.N. and J.R.C. wrote the manuscript.

## Acknowledgments

This work was funded by a research grant to J.R.C. from the Spanish Ministry of Science and Innovation (PID2023-146143NB-I00). It benefitted also from an institutional grant from the Severo Ochoa Program for Centers of Excellence in R&D (CEX2023-001386-S/AEI/10.13039/501100011033). We thank the Diamond Light Source for Titan Krios data collection at the UK national electron Bio-Imaging Center (eBIC) under proposal BI22006.

## Suplementary information

**Figure S1.**
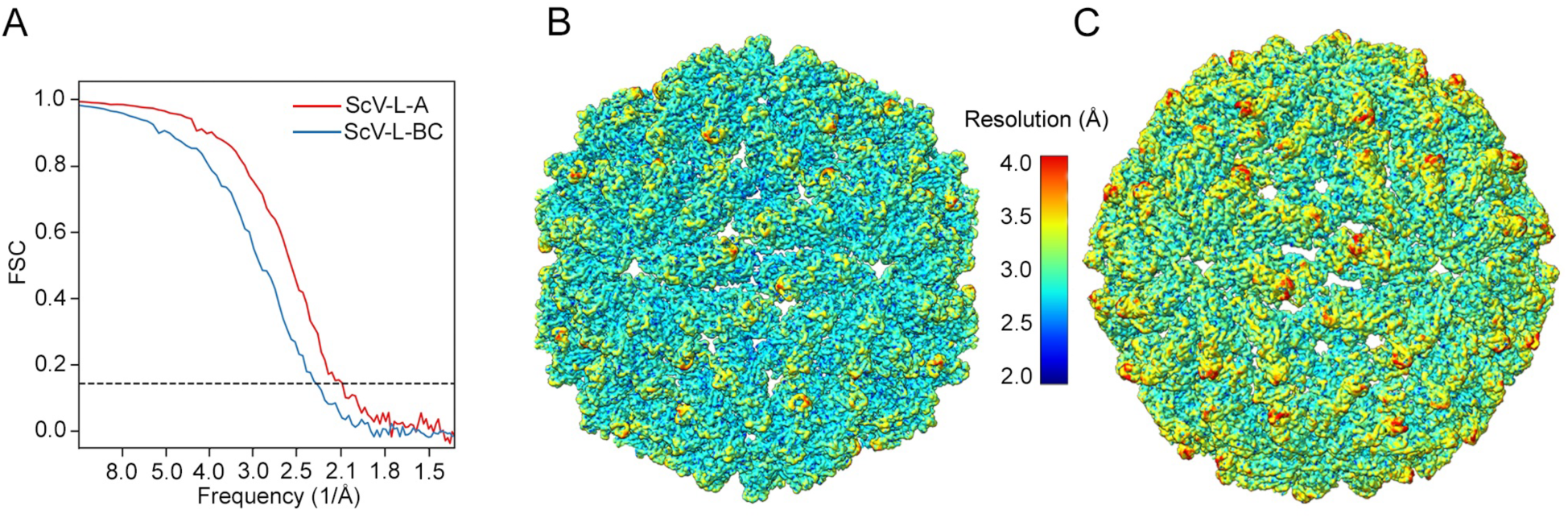
Fourier shell correlation analysis of the ScV-L-A and ScV-L-BC cryo-EM reconstructions. (A) Gold-standard Fourier shell correlation (FSC) curves for the ScV-L-A (red) and ScV-L-BC (blue) reconstructions. Independent half-maps were refined separately, and the nominal resolution of each reconstruction was estimated using the FSC = 0.143 criterion (dashed line). (B, C) Local-resolution maps of ScV-L-A (B) and ScV-L-BC (C) colored according to local resolution (Å; color scale), showing the distribution of resolution across the capsid densities.

**Figure S2.**
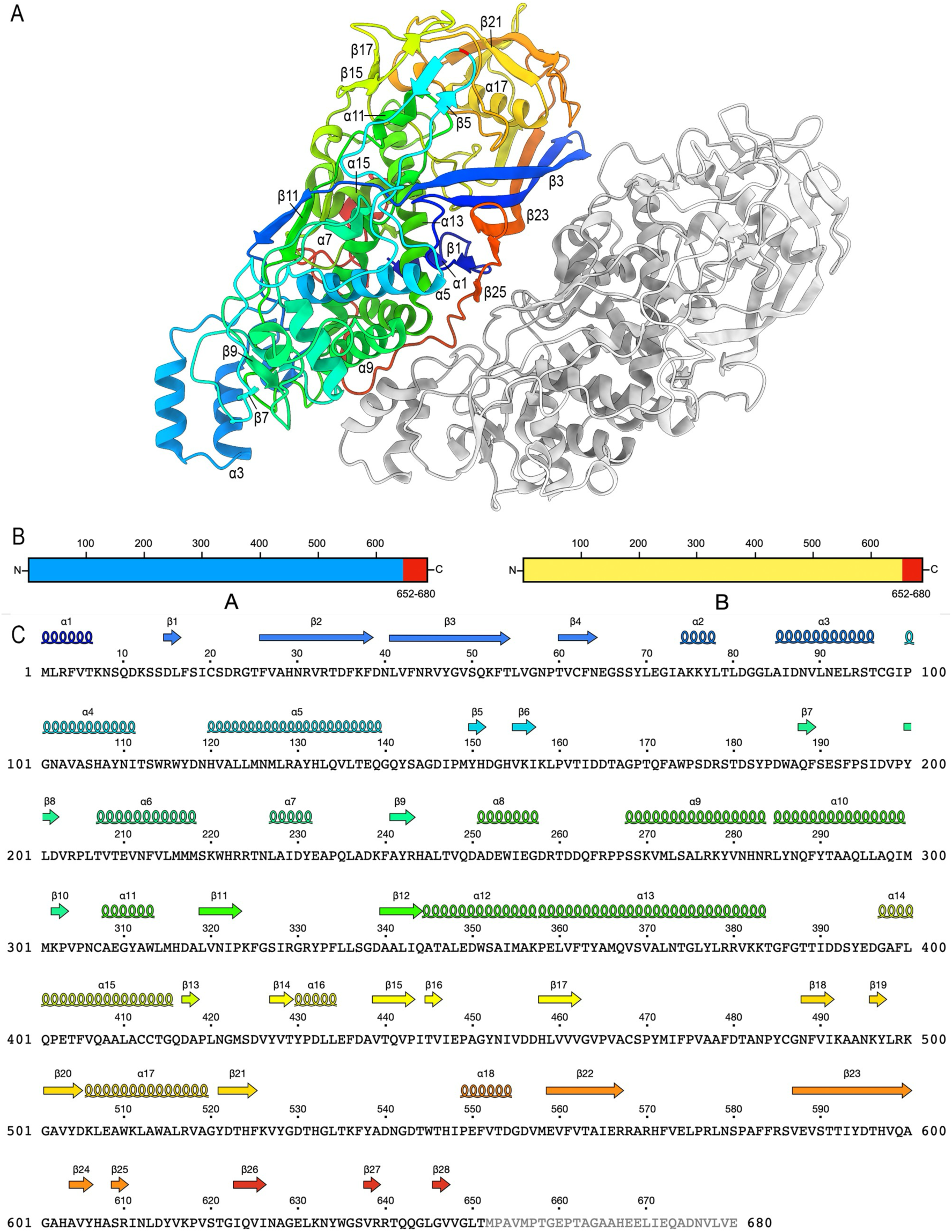
Atomic model and secondary-structure organization of the ScV-L-A Gag capsid protein. (A) Ribbon diagram of the asymmetric unit of the ScV-L-A capsid. The A subunit is rainbow-colored from blue (N terminus) to red (C terminus); the B subunit is in gray. Secondary-structure elements are labelled. (B) Schematic representation of the modeled regions of Gag in conformations A (blue) and B (yellow), highlighting the unresolved C-terminal region (Met652–Glu680) in red. (C) Gag sequence and corresponding secondary-structure elements with the same color scheme used in panel A (spirals, α-helices; arrows, β-strands). The unresolved C-terminal residues are shown in grey.

**Figure S3.**
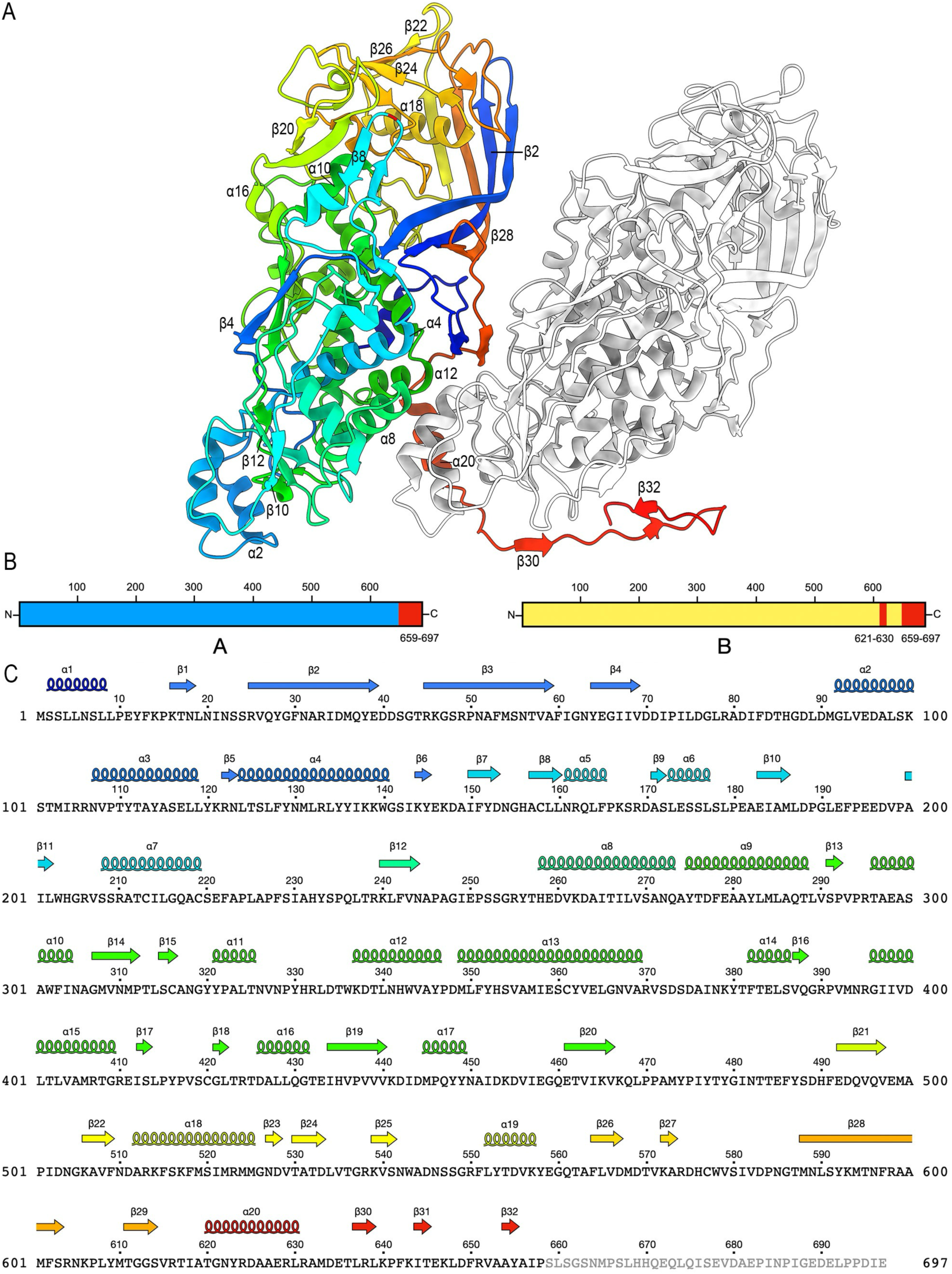
Atomic model and secondary-structure organization of the ScV-L-BC Gag capsid protein. (A) Ribbon diagram of the asymmetric unit of the ScV-L-BC capsid. The A subunit is rainbow-colored from blue (N terminus) to red (C terminus); the B subunit is in gray. Secondary-structure elements are labelled. (B) Schematic representation of the modeled regions of Gag in conformations A (blue) and B (yellow), highlighting the unresolved regions in red. The internal unresolved segment Gly621–Leu630 of CP-B is indicated. (C) Gag sequence and corresponding secondary-structure elements with the same color scheme used in panel A (spirals, α-helices; arrows, β-strands). The unresolved C-terminal residues are shown in grey.

**Figure S4.**
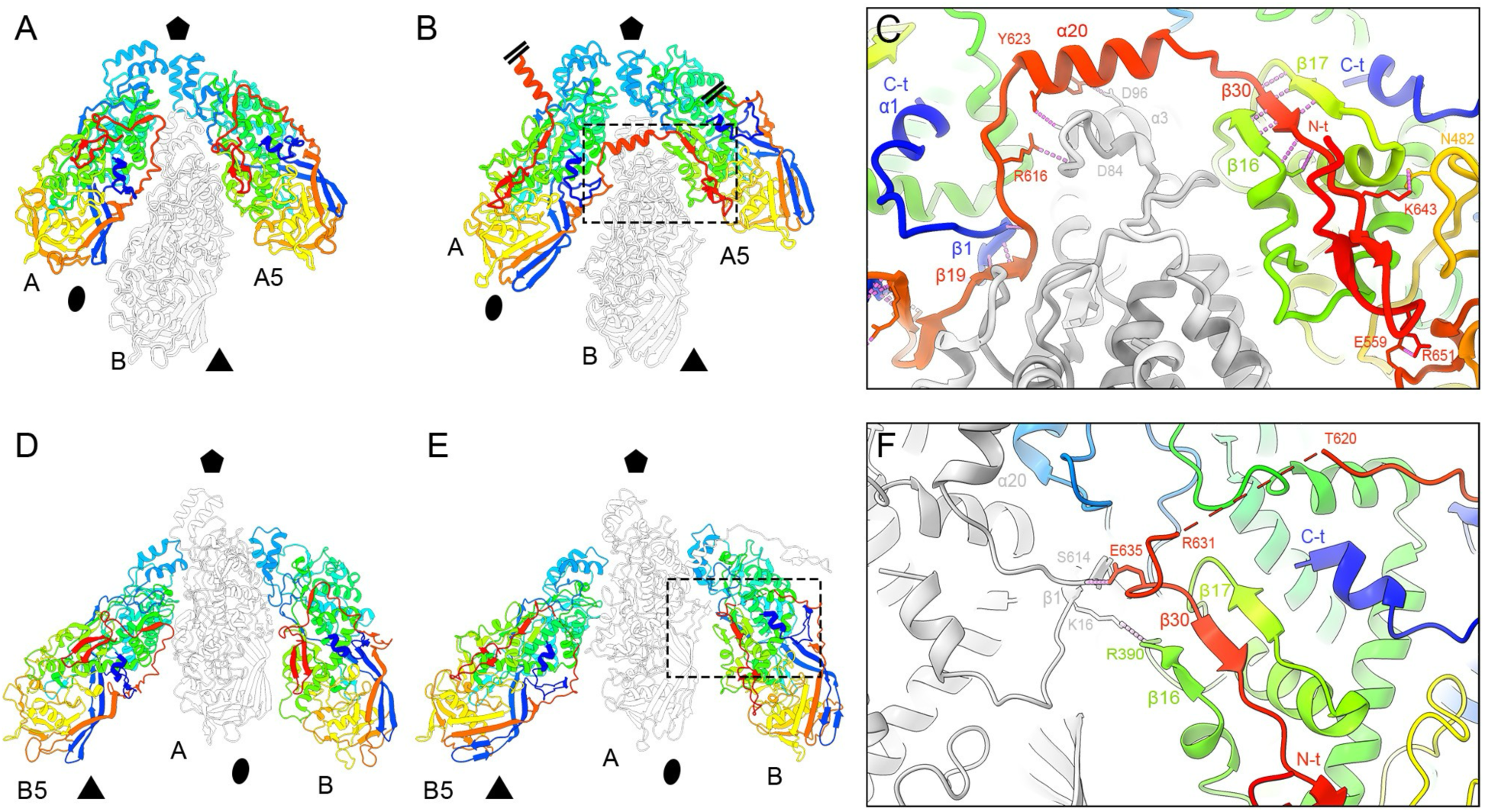
β-strand augmentation and molecular swapping between Gag subunits in ScV-L-BC. (A, B) Comparison of three Gag subunits within the same decamer: two CP-A conformers (rainbow gradient, A and A5) and one CP-B conformer (grey) in ScV-L-A (A) and ScV-L-BC (B). The extended C-terminal region of ScV-L-BC CP-A mediates molecular swapping with the adjacent CP-A subunit. (C) Closeup view of the β-strand augmentation between neighbouring CP-A subunits in ScV-L-BC. The C-terminal β30 strand from one CP-A subunit inserts between β16 and β17 of the adjacent CP-A subunit, forming an intermolecular β-sheet. (D, E) Equivalent comparison involving two CP-B conformers (rainbow gradient, B5 and B) and one CP-A conformer (grey) in ScV-L-A (D) and ScV-L-BC (E). (F) Closeup view of the β16–β17–β30 sheet formed within the ScV-L-BC CP-B conformer through intramolecular interactions. The unresolved Gly621–Leu630 segment in CP-B (dashed line) is indicated.

**Table S1.** Cryo-EM data collection and refinement statistics for ScV-L-A and ScV-L-BC.

|  | ScV-L-A<br>PDB: 9rk2<br>EMD-54013 | ScV-L-BC<br>PDB: 9suj<br>EMD-55221 |
| --- | --- | --- |
| **Data collection and processing** |  |  |
| Microscope | Titan Krios | Titan Krios |
| Detector | Gatan K3 | Gatan K3 |
| Magnification | 81,000x | 81,000x |
| Voltage (kV) | 300 | 300 |
| Electron exposure (e<sup>-</sup>/Å<sup>2</sup>) | 31.2 | 31.2 |
| Exposure per frame (e<sup>-</sup>/Å<sup>2</sup>) | 1.04 | 1.04 |
| Defocus range (μm) | -0.8 to -2.3 | -0.8 to -2.3 |
| Pixel size (Å) | 1.06 | 1.06 |
| Micrographs collected (no.) | 12248 | 12248 |
| Initial particles (no.) | 469,633 | 469,633 |
| Final particles (no.) | 280,585 | 29,699 |
| Symmetry imposed | I2 | I2 |
| Map resolution (Å) | 2.1 | 2.3 |
| FSC threshold | 0.143 | 0.143 |
| **Refinement** |  |  |
| Mask correlation coefficient | 0.83 | 0.83 |
| Map sharpening software | Cryosparc | Cryosparc |
| **Model composition** |  |  |
| Non-hydrogenatoms | 618,120 | 619,800 |
| Protein residues | 78,120 | 78,360 |
| **ADP (B-factors)** |  |  |
| min | 6.50 | 0.00 |
| max | 112.17 | 106.52 |
| mean | 32.62 | 34.55 |
| **R.m.s deviations** |  |  |
| Bond lengths (Å) | 0.006 | 0.005 |
| Bond angles (°) | 0.962 | 0.966 |
| **Validation** |  |  |
| Molprobity score | 1.92 | 1.58 |
| Clashscore | 2.77 | 3.95 |
| Rotamer outliers (%) | 4.68 | 2.48 |
| **Ramachandran plot** |  |  |
| Favored (%) | 95.00 | 97.54 |
| Allowed (%) | 4.92 | 2.46 |
| Outliers (%) | 0.08 | 0.00 |

## REFERENCES

1. Mu F, Li B, Cheng S, Jia J, Jiang D, Fu Y, et al. 2021. Nine viruses from eight lineages exhibiting new evolutionary modes that co-infect a hypovirulent phytopathogenic fungus. PLoS Pathog 17:e1009823. 10.1371/journal.ppat.1009823.

2. Thapa V, Roossinck MJ. 2019. Determinants of Coinfection in the Mycoviruses. Front Cell Infect Microbiol 9. 10.3389/fcimb.2019.00169.

3. Poimala A, Parikka P, Hantula J, Vainio EJ. 2021. Viral diversity in Phytophthora cactorum population infecting strawberry. Environ Microbiol 23:5200–21. 10.1111/1462-2920.15519.

4. Duan J, Yao Y, Xu J, Zhang A, Kong X, Lin Y, et al. 2025. The rules in co-infection of multiple viruses across diverse lineages in a fungal host. mBio 16:e00262–25. 10.1128/mbio.00262-25.

5. Hollings M. 1962. Viruses Associated with A Die-Back Disease of Cultivated Mushroom. Nature 196:4858. 10.1038/196962a0.

6. Hough B, Steenkamp E, Wingfield B, Read D. 2023. Fungal Viruses Unveiled: A Comprehensive Review of Mycoviruses. Viruses 15:1202. 10.3390/v15051202.

7. Yu X, Li B, Fu Y, Xie J, Cheng J, Ghabrial SA, et al. 2013. Extracellular transmission of a DNA mycovirus and its use as a natural fungicide. Proc Natl Acad Sci U S A 110:1452–7. 10.1073/pnas.1213755110.

8. Buivydaite Z, Winding A, Sapkota R. 2024. Transmission of mycoviruses: new possibilities. Front Microbiol 15. 10.3389/fmicb.2024.1432840.

9. Xie J, Jiang D. 2024. Understanding the Diversity, Evolution, Ecology, and Applications of Mycoviruses. Annu Rev Microbiol 78:595–620. 10.1146/annurev-micro-041522-105358.

10. Ayllón MA, Vainio EJ. 2023. Mycoviruses as a part of the global virome: Diversity, evolutionary links and lifestyle. Adv Virus Res 115:1–86. 10.1016/bs.aivir.2023.02.002.

11. Villan Larios DC, Diaz Reyes BM, Pirovani CP, Loguercio LL, Santos VC, Góes-Neto A, et al. 2023. Exploring the Mycovirus Universe: Identification, Diversity, and Biotechnological Applications. J Fungi 9:361. 10.3390/jof9030361.

12. Lin YH, Fujita M, Chiba S, Hyodo K, Andika IB, Suzuki N, et al. 2019. Two novel fungal negative-strand RNA viruses related to mymonaviruses and phenuiviruses in the shiitake mushroom (Lentinula edodes). Virology 533:125–36. 10.1016/j.virol.2019.05.008.

13. Dai R, Yang S, Pang T, Tian M, Wang H, Zhang D, et al. 2024. Identification of a negative-strand RNA virus with natural plant and fungal hosts. Proc Natl Acad Sci U S A 121:e2319582121. 10.1073/pnas.2319582121.

14. Marzano SYL, Nelson BD, Ajayi-Oyetunde O, Bradley CA, Hughes TJ, Hartman GL, et al. 2016. Identification of Diverse Mycoviruses through Metatranscriptomics Characterization of the Viromes of Five Major Fungal Plant Pathogens. J Virol 90:6846–63. 10.1128/JVI.00357-16.

15. Donaire L, Ayllón MA. 2017. Deep sequencing of mycovirus-derived small RNAs from Botrytis species. Mol Plant Pathol 18:1127–37. 10.1111/mpp.12466.

16. Hillman BI, Suzuki N. 2004. Viruses of the chestnut blight fungus, Cryphonectria parasitica. Adv Virus Res 63:423–72. 10.1016/S0065-3527(04)63007-7.

17. Dálya LB, Černý M, de la Peña M, Poimala A, Vainio EJ, Hantula J, et al. 2024. Diversity and impact of single-stranded RNA viruses in Czech Heterobasidion populations. mSystems 9:e0050624. 10.1128/msystems.00506-24.

18. Li P, Bhattacharjee P, Wang S, Zhang L, Ahmed I, Guo L. 2019. Mycoviruses in Fusarium Species: An Update. Front Cell Infect Microbiol 9:257. 10.3389/fcimb.2019.00257.

19. Illana A, Marconi M, Rodríguez-Romero J, Xu P, Dalmay T, Wilkinson MD, et al. 2017. Molecular characterization of a novel ssRNA ourmia-like virus from the rice blast fungus Magnaporthe oryzae. Arch Virol 162:891–5. 10.1007/s00705-016-3144-9.

20. Liu L, Xie J, Cheng J, Jiang D. 2014. Fungal negative-stranded RNA virus that is related to bornaviruses and nyaviruses. Proc Natl Acad Sci U S A 111:12205–10. 10.1073/pnas.1401786111.

21. Zhang R, Hisano S, Tani A, Kondo H, Kanematsu S, Suzuki N. 2016. A capsidless ssRNA virus hosted by an unrelated dsRNA virus. Nat Microbiol 1:1. 10.1038/nmicrobiol.2015.1.

22. Fadli M, Hisano S, Novoa G, Castón JR, Kondo H, Suzuki N. 2025. A capsidless (+)RNA yadokarivirus hosted by a dsRNA virus is infectious as particles, cDNA, and dsRNA. J Virol 99:e02166–24. 10.1128/jvi.02166-24.

23. Naitow H, Tang J, Canady M, Wickner RB, Johnson JE. 2002. L-A virus at 3.4 Å resolution reveals particle architecture and mRNA decapping mechanism. Nat Struct Mol Biol 9:10. 10.1038/nsb844.

24. Grybchuk D, Procházková M, Füzik T, Konovalovas A, Serva S, Yurchenko V, et al. 2022. Structures of L-BC virus and its open particle provide insight into Totivirus capsid assembly. Commun Biol 5:1. 10.1038/s42003-022-03793-z.

25. Pan J, Dong L, Lin L, Ochoa WF, Sinkovits RS, Havens WM, et al. 2009. Atomic structure reveals the unique capsid organization of a dsRNA virus. Proc Natl Acad Sci U S A 106:4225–30. 10.1073/pnas.0812071106.

26. Luque D, Gómez-Blanco J, Garriga D, Brilot AF, González JM, Havens WM, et al. 2014. Cryo-EM near-atomic structure of a dsRNA fungal virus shows ancient structural motifs preserved in the dsRNA viral lineage. Proc Natl Acad Sci U S A 111:7641–6. 10.1073/pnas.1404330111.

27. Mata CP, Luque D, Gómez-Blanco J, Rodríguez JM, González JM, Suzuki N, et al. 2017. Acquisition of functions on the outer capsid surface during evolution of double-stranded RNA fungal viruses. PLoS Pathog 13:e1006755. 10.1371/journal.ppat.1006755.

28. Wang H, Salaipeth L, Miyazaki N, Suzuki N, Okamoto K. 2023. Capsid structure of a fungal dsRNA megabirnavirus reveals its previously unidentified surface architecture. PLoS Pathog 19:e1011162. 10.1371/journal.ppat.1011162.

29. Wang H, Marucci G, Munke A, Hassan MM, Lalle M, Okamoto K. 2024. High-resolution comparative atomic structures of two Giardiavirus prototypes infecting G. duodenalis parasite. PLoS Pathog 20:e1012140. 10.1371/journal.ppat.1012140.

30. Okamoto K, Miyazaki N, Larsson DSD, Kobayashi D, Svenda M, Mühlig K, et al. 2016. The infectious particle of insect-borne totivirus-like Omono River virus has raised ridges and lacks fibre complexes. Sci Rep 6:33170. 10.1038/srep33170.

31. Cheng Y. 2015. Single-Particle Cryo-EM at Crystallographic Resolution. Cell 161:450–457. 10.1016/j.cell.2015.03.049.

32. Wickner RB, Fujimura T, Esteban R. 2013. Viruses and prions of Saccharomyces cerevisiae. Adv Virus Res 86:1–36. 10.1016/B978-0-12-394315-6.00001-5.

33. Taggart NT, Crabtree AM, Creagh JW, Bizarria RJ, Li S, de la Higuera I, et al. 2023. Novel viruses of the family Partitiviridae discovered in Saccharomyces cerevisiae. PLoS Pathog 19:e1011418. 10.1371/journal.ppat.1011418.

34. Vijayraghavan S, Kozmin SG, Xi W, McCusker JH. 2023. A novel narnavirus is widespread in Saccharomyces cerevisiae and impacts multiple host phenotypes. G3 (Bethesda) 13:jkac337. 10.1093/g3journal/jkac337.

35. Dinman JD, Icho T, Wickner RB. 1991. A -1 ribosomal frameshift in a double-stranded RNA virus of yeast forms a gag-pol fusion protein. Proc Natl Acad Sci U S A 88:174–8. 10.1073/pnas.88.1.174.

36. Vijayraghavan S, Kozmin SG, Strope PK, Skelly DA, Magwene PM, Dietrich FS, et al. 2023. RNA viruses, M satellites, chromosomal killer genes, and killer/nonkiller phenotypes in the 100-genomes S. cerevisiae strains. G3 (Bethesda) 13:jkad167. 10.1093/g3journal/jkad167.

37. Schmidt L, Tüting C, Kyrilis FL, Hamdi F, Semchonok DA, Hause G, et al. 2024. Delineating organizational principles of the endogenous L-A virus by cryo-EM and computational analysis of native cell extracts. Commun Biol 7:557. 10.1038/s42003-024-06204-7.

38. Fujimura T, Esteban R. 2013. Cap Snatching in Yeast L-BC Double-stranded RNA Totivirus. J Biol Chem 288:23716–24. 10.1074/jbc.M113.490953.

39. Chau S, Gao J, Diao AJ, Cao SB, Azhieh A, Davidson AR, et al. 2023. Diverse yeast antiviral systems prevent lethal pathogenesis caused by the L-A mycovirus. Proc Natl Acad Sci U S A 120:e2208695120. 10.1073/pnas.2208695120.

40. Cheng RH, Caston JR, Wang GJ, Gu F, Smith TJ, Baker TS, et al. 1994. Fungal virus capsids, cytoplasmic compartments for the replication of double-stranded RNA, formed as icosahedral shells of asymmetric Gag dimers. J Mol Biol 244:255–8. 10.1006/jmbi.1994.1726.

41. Fujimura T, Esteban R. 2012. Cap Snatching of Yeast L-A Double-stranded RNA Virus Can Operate in Trans and Requires Viral Polymerase Actively Engaging in Transcription. J Biol Chem 287:12797–804. 10.1074/jbc.M111.327676.

42. Duquerroy S, Da Costa B, Henry C, Vigouroux A, Libersou S, Lepault J, et al. 2009. The picobirnavirus crystal structure provides functional insights into virion assembly and cell entry. EMBO J 28:1655–65. 10.1038/emboj.2009.109.

43. Procházková M, Füzik T, Grybchuk D, Yurchenko V, Plevka P. 2022. Virion structure of Leishmania RNA virus 1. Virology 577:149–54. 10.1016/j.virol.2022.09.014.

44. Stevens A, Muratore K, Cui Y, Johnson PJ, Zhou ZH. 2021. Atomic Structure of the Trichomonas vaginalis Double-Stranded RNA Virus 2. mBio 12:e02924–20. 10.1128/mBio.02924-20.

45. Shao Q, Jia X, Gao Y, Liu Z, Zhang H, Tan Q, et al. 2021. Cryo-EM reveals a previously unrecognized structural protein of a dsRNA virus implicated in its extracellular transmission. PLoS Pathog 17:e1009396. 10.1371/journal.ppat.1009396.

46. Dunn SE, Li H, Cardone G, Nibert ML, Ghabrial SA, Baker TS. 2013. Three-dimensional Structure of Victorivirus HvV190S Suggests Coat Proteins in Most Totiviruses Share a Conserved Core. PLoS Pathog 9:e1003225. 10.1371/journal.ppat.1003225.

47. Tang J, Ochoa WF, Sinkovits RS, Poulos BT, Ghabrial SA, Lightner DV, et al. 2008. Infectious myonecrosis virus has a totivirus-like, 120-subunit capsid, but with fiber complexes at the fivefold axes. Proc Natl Acad Sci U S A 105:17526–31. 10.1073/pnas.0806724105.

48. Celitan E, Stanevičienė R, Servienė E, Serva S. 2024. Highly stable Saccharomyces cerevisiae L-BC capsids with versatile packing potential. Front Bioeng Biotechnol 12. 10.3389/fbioe.2024.1456453.

49. Rohou A, Grigorieff N. 2015. CTFFIND4: Fast and accurate defocus estimation from electron micrographs. J Struct Biol 192:216–21. 10.1016/j.jsb.2015.08.008.

50. de la Rosa-Trevín JM, Otón J, Marabini R, Zaldívar A, Vargas J, Carazo JM, et al. 2013. Xmipp 3.0: an improved software suite for image processing in electron microscopy. J Struct Biol 184:321–8. 10.1016/j.jsb.2013.09.015.

51. Punjani A, Rubinstein JL, Fleet DJ, Brubaker MA. 2017. cryoSPARC: algorithms for rapid unsupervised cryo-EM structure determination. Nat Methods 14:290–6. 10.1038/nmeth.4169.

52. Vilas JL, Gómez-Blanco J, Conesa P, Melero R, Miguel de la Rosa-Trevín J, Otón J, et al. 2018. MonoRes: Automatic and Accurate Estimation of Local Resolution for Electron Microscopy Maps. Structure 26:337–344.e4. 10.1016/j.str.2017.12.018.

53. Liebschner D, Afonine PV, Baker ML, Bunkóczi G, Chen VB, Croll TI, et al. 2019. Macromolecular structure determination using X-rays, neutrons and electrons: recent developments in Phenix. Acta Crystallogr D Struct Biol 75:861–77. 10.1107/S2059798319011471.

54. Emsley P, Lohkamp B, Scott WG, Cowtan K. 2010. Features and development of Coot. Acta Crystallogr D Biol Crystallogr 66:486–501. 10.1107/S0907444910007493.

55. Williams CJ, Headd JJ, Moriarty NW, Prisant MG, Videau LL, Deis LN, et al. 2018. MolProbity: More and better reference data for improved all-atom structure validation. Protein Sci 27:293–315. 10.1002/pro.3330.

56. Meng EC, Goddard TD, Pettersen EF, Couch GS, Pearson ZJ, Morris JH, et al. 2023. UCSF ChimeraX: Tools for structure building and analysis. Protein Sci 32:e4792. 10.1002/pro.4792.

57. Holm L. 2022. Dali server: structural unification of protein families. Nucleic Acids Res 50:W210–5. 10.1093/nar/gkac387.

